# A three-dimensional culture model of innervated human skeletal muscle enables studies of the adult neuromuscular junction and disease modeling

**DOI:** 10.1101/275545

**Authors:** Mohsen Afshar Bakooshli, Ethan S Lippmann, Ben Mulcahy, Nisha R Iyer, Christine T Nguyen, Kayee Tung, Bryan A Stewart, Hubrecht van den Dorpel, Tobias Fuehrmann, Molly S Shoichet, Anne Bigot, Elena Pegoraro, Henry Ahn, Howard Ginsberg, Mei Zhen, Randolph S Ashton, Penney M Gilbert

## Abstract

Two-dimensional (2D) human skeletal muscle fiber cultures are ill equipped to support the contractile properties of maturing muscle fibers. This limits their application to the study of adult human neuromuscular junction (NMJ) development, a process requiring maturation of muscle fibers in the presence of motor neuron endplates. Here we describe a three-dimensional (3D) co-culture method whereby human muscle progenitors mixed with human pluripotent stem cell-derived motor neurons self-organize to form functional NMJ connections within two weeks. Functional connectivity between motor neuron endplates and muscle fibers is confirmed with calcium transient imaging and electrophysiological recordings. Notably, we only observed epsilon acetylcholine receptor subunit protein upregulation and activity in 3D co-culture. This demonstrates that the 3D co-culture system supports a developmental shift from the embryonic to adult form of the receptor that does not occur in 2D co-culture. Further, 3D co-culture treatments with myasthenia gravis patient sera shows the ease of studying human disease with the system. This work delivers a simple, reproducible, and adaptable method to model and evaluate adult human NMJ de novo development and disease in culture.

## Introduction

The skeletal muscle neuromuscular junction (NMJ) is a highly organized synapse formed between a motor neuron (MN) axon and a muscle fiber. It is designed to transmit efferent signals from projecting MNs to muscle fibers in order to actuate fiber contraction. Nicotinic acetylcholine receptors (AChRs) clustered at the NMJ’s postsynaptic muscle fiber membrane mediate this signal by binding acetylcholine (ACh) neurotransmitters released from vesicles at the presynaptic MN axon terminal. AChRs are ligand-gated ion channels composed of five protein subunits. During development the gamma subunit in embryonic AChRs is replaced by an epsilon subunit in the adult synapse (Mishina et al., 1986; Missias et al., 1996). Previous animal studies showed that this AChR subunit transition occurs in the presence of motor axon endplates and confirmed that transcription of the epsilon gene (CHRNE) is stimulated by AChR Inducing Activity (ARIA) via ErbB receptors, a nerve derived ligand of the neuregulin-1 (NRG1) family (Martinou et al., 1991). Consistently, CHRNE transcripts are detected in rodent 2D and 3D skeletal muscle fiber cultures when co-cultured with nerve cells (Bach et al., 2003; Ostrovidov et al., 2017; Smith et al., 2016; Vilmont et al., 2016). However, despite significant progress toward directing human pluripotent stem cells (PSCs) to the motor neuron lineage (Ashton et al., 2015; Hu and Zhang, 2010; Lippmann et al., 2014; Maury et al., 2015; Shimojo et al., 2015; Zhang et al., 2001) and establishing electrically and chemically responsive human muscle fibers in vitro (Madden et al., 2015), the first reports of human NMJ models – 2D human muscle fiber and motor neuron co-cultures – were unable to demonstrate synapse maturation via the gamma to epsilon AChR subunit switch (Guo et al., 2011; Steinbeck et al., 2016), and there are no reports of epsilon AChR protein expression or function in culture in the absence of enforced gene expression.

Congenital myasthenic syndrome is one of the most prevalent genetic diseases of the NMJ and commonly arises from mutations in one of the AChR encoding genes (Engel et al., 2010). The vast majority of mutations causing the disease arise in the CHRNE gene, the adult specific subunit of the AChR (Abicht et al., 2012; Engel et al., 1993). Given the lack of effective therapies for a wide range of neuromuscular diseases impacting the adult NMJ (Ohno et al., 1999), and that the majority of AChR mutations are mutations of the CHRNE gene (Ohno et al., 1995), a robust method to model the adult human NMJ in a dish is needed to synergize with recent advances in differentiating patient-derived PSCs to the MN lineage (Chen et al., 2011; Hu et al., 2010; Lorenz et al., 2017; Sances et al., 2016).

Here we report a method integrating architectural cues with co-culture techniques to create an environment conducive to formation of the adult human NMJ in as early as two weeks. We show that a 3D culture system is essential for long-term maintenance of maturing muscle fibers in culture. It supports the formation and morphological maturation of AChR clusters primed for synaptogenesis and the de novo transition from the embryonic to the adult NMJ composition upon contact with MN endplates. We demonstrate that agrin, a trophic factor produced by MNs, induces AChR clustering and maturation in 3D muscle fiber cultures. We confirm formation of functional NMJ connections by imaging muscle fiber calcium transients and capturing electrophysiological recordings in response to glutamate-induced MN firing and that treatment with inhibitors targeting pre- and post-synapse function block this firing. We show that the 3D co-culture platform, and not a 2D co-culture system, supports the transition from the embryonic to the adult AChR, thereby enabling the functional assessment of the adult neuromuscular junction in vitro. Our data align with prior studies showing that epsilon functional activity is regulated post-transcriptionally (Bruneau et al., 2005; Caroni et al., 1993; Jayawickreme and Claudio, 1994; Khan et al., 2014; Missias et al., 1996; Ross et al., 1991; Wild et al., 2016; Witzemann et al., 2013; Xu and Salpeter, 1997; Yampolsky et al., 2008), and in particular, supports work indicating a role for innervation (spontaneous miniature endplate potentials) and / or muscle fiber maturation in encouraging subunit substitution (Caroni et al., 1993; Missias et al., 1996; Witzemann et al., 2013; Xu and Salpeter, 1997; Yampolsky et al., 2008). Finally, we demonstrate the versatility and ease of our system for modelling human disease by treating neuromuscular co-cultures with IgG purified from myasthenia gravis (MG) patient sera together with human complement, which results in readily visible clinical phenotypes in as early as two weeks of culture time. Thus, the described 3D co-culture model enables investigation of adult human NMJ development and therefore adult forms of neuromuscular diseases in vitro for the first time.

## Results

### Myogenic differentiation in 3D enhances fiber maturation and AChR clustering

We performed a side-by-side comparison of human skeletal muscle fiber populations derived in standard 2D culture versus 3D culture and uncovered differences in fiber maturation and AChR clustering (**Figure S1A**). We established primary myogenic progenitor and fibroblast-like cell lines from human biopsy tissues (Blau and Webster, 1981) (**Figure S1B**), and seeded them at defined ratios either within a fibrin/Geltrex™ hydrogel (3D) or into 12-well tissue culture plastic dishes coated with Geltrex™ (2D) or a fibrinogen/Geltrex™ blend (**Figure S1A**). Muscle cell laden hydrogels were formed within a polydimethylsiloxane channel and anchored at each end of the channel to the nylon hooks of Velcro™ fabric, which act as artificial tendons and establish uniaxial tension during 3D tissue remodelling and differentiation (Bell et al., 1979; Madden et al., 2015; Vandenburgh et al., 1988) (**Figure S1C**).

Immunofluorescence analysis of the muscle contractile protein sarcomeric a-actinin (SAA) revealed the uniform alignment of striated muscle fibers along the tension axis in the 3D tissues (**Figure 1A** and **Figure S1E**), while 2D muscle fiber cultures were regionally aligned (**Figure 1A**), but globally disorganized (**Figure S1D**). In contrast to the muscle fibers established in 2D cultures, those derived in 3D culture progressively increased in diameter over three weeks in culture (**Figure 1B**) while maintaining fiber alignment and assembled contractile apparatus (**Figure 1A**). Furthermore, over time in 3D culture, muscle tissues upregulated expression of the fast and slow adult isoforms of myosin heavy chain (MHC), which was accompanied by a downregulation of embryonic MHC expression, suggesting a gradual sarcomere structural maturation (**Figure 1C** and **Figures S2A-E**). The absence of these trends in 2D muscle fiber culture may be explained by the inability of tissue culture plastic to support muscle fiber contraction resulting in the increased incidence of damaged fibers observed in 2D cultures (**Figure S1D**) and an enrichment of small, immature fibers (**Figures 1A-B** and **Figures S2G-H**).

**Figure 1.**
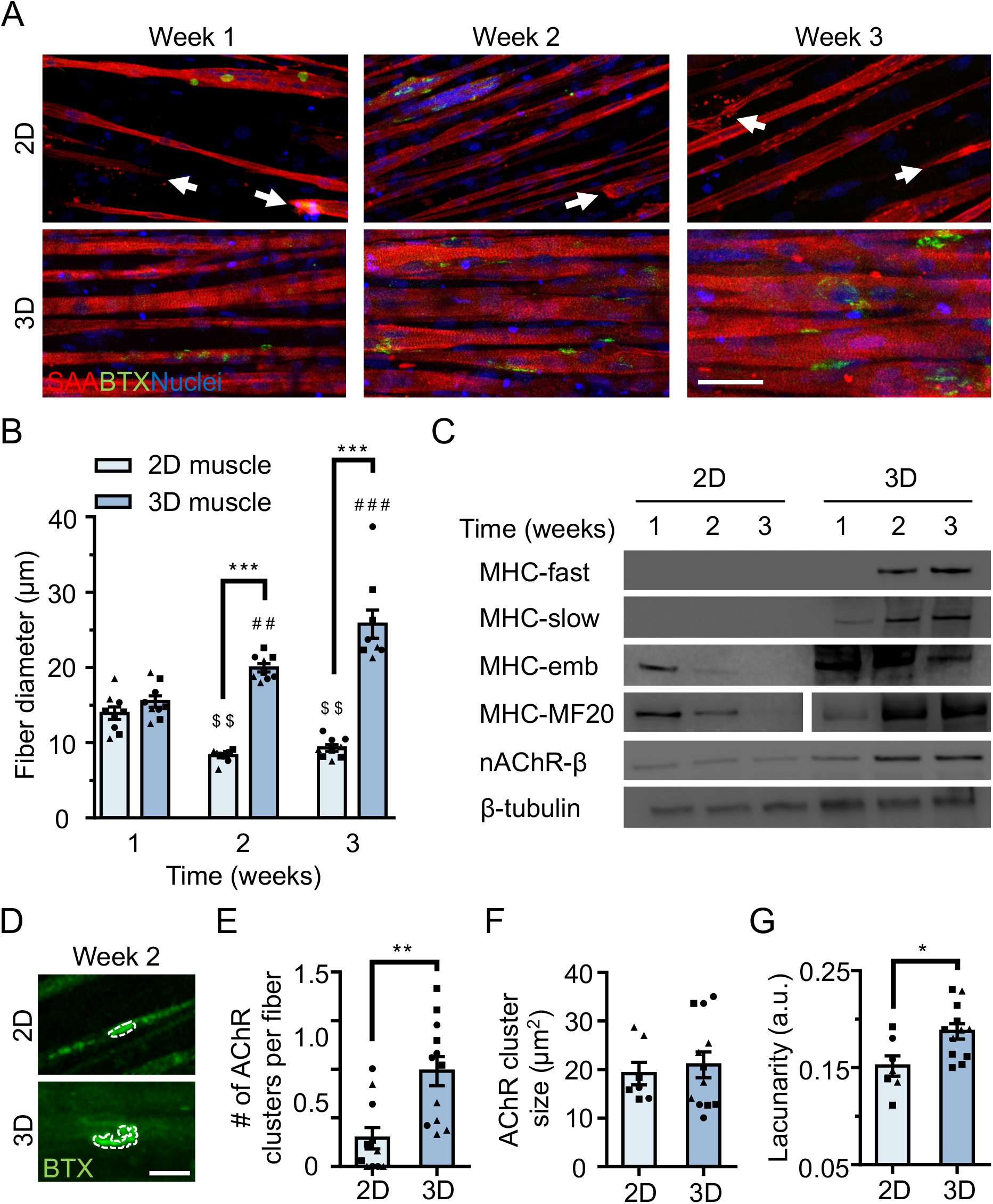
3D culture enhances skeletal muscle fiber maturation over 2D culture. **(A)** Representative confocal images of muscle fibers established in 2D (top row) and 3D conditions and immunostained for sarcomeric a-actinin (SAA; red), a-bungarotoxin (BTX; green), and Hoechst 33342 (blue) after 1, 2, and 3 weeks of culture. Scale bar, 50 μm. White arrowheads indicate broken fibers. **(B)** Bar graph of muscle fiber diameter quantified in 2D (light blue) and 3D (blue) cultures over time. n = 9 independent samples from 3 muscle patient donors. A minimum of 50 myotubes per time point per patient sample were analyzed. ^##^p < 0.01 and ^###^p < 0.001 compared with 3D cultures at week 1. ^$$^p < 0.01 compared with 2D culture at week 1. **(C)** Representative western blot images of myosin heavy chain (MHC) isoforms (fast, slow, embryonic (emb), and pan (MF-20)) nicotinic AChR-β (nAChR-β), and β-tubulin in 2D compared with 3D cultures over time. **(D)** Representative confocal images of muscle fibers cultured in 2D or 3D for two weeks and then labeled with a-bungarotoxin (green). AChR clusters are outlined with white dashed lines. Scale bar, 25 μm. **(E-G)** Bar graphs indicating average **(E)** number of AChR clusters per fiber, **(F)** AChR cluster size, and **(G)** AChR cluster lacunarity in 2D (light blue) and 3D (blue) muscle fiber cultures at week 2. n = minimum of 9 independent samples from 3 muscle patient donors. A minimum of 30 microscopic images per culture condition were analyzed. In (B), (C), and (E-G) each symbol represents data from one muscle patient donor. Values in (B), (E), (F), and (G) are mean ± SEM. * p < 0.05, ** p < 0.01, *** p < 0.001.

In support of our molecular characterization, 3D human muscle tissues were capable of generating active force in as early as 10 days of differentiation as evidenced by spontaneous twitches (**Movie S1**), which were not observed in 2D cultures. Consistent with prior reports (Madden et al., 2015), two-week old 3D muscle tissue twitch response could be paced by low frequency electrical stimuli (1 Hz; **Movie S1**), which converted into tetanus contractions in response to increased frequency (20 Hz; **Movie S1**). Similarly, ACh stimulation (10 μM) produced an immediate tetanus response (**Movie S1**) in 3D tissues suggesting an abundance of active AChRs, while the response of 2D muscle fiber cultures at this time-point was significantly less and inevitably resulted in muscle fiber damage and / or release from the culture substrate (**Movie S2**).

To evaluate the calcium handling capacity of 3D muscle fiber cultures, we transduced human muscle progenitor cells with lentiviral particles encoding GCaMP6 (Chen et al., 2013), a sensitive calcium indicator protein, driven by the MHCK7 (Madden et al., 2015) promoter, a muscle specific gene. Muscle fibers in 3D tissues generated strong collective calcium transient in response to electrical stimulation and immediately following exposure to ACh (**Figures S3A-C** and **Movie S3**).

To evaluate the electrophysiological characteristics of single muscle fibers in 3D cultures, muscle progenitor cells were stably transduced with a light-gated ion channel, channelrhodopsin-2 (ChR2), driven by an EF1a promoter (Zhang et al., 2007). 3D muscle tissues generated using optogenetically-responsive muscle progenitor cells contracted in response to light stimulation on the second week of the culture (**Movie S4**). Next, single muscle fiber membrane potentials were recorded in these tissues using sharp microelectrode recording (**Figure S3D**). As expected, recordings of 3D muscle prior to light stimulation revealed little electrical activity (**Figure S3E**), while light activation generated a clear depolarization of the membrane potential (**Figure S3F**). We also took a more traditional approach using single, sharp electrode electrophysiology to measure membrane potential and test excitability. Passing depolarizing current led to regenerative potentials that become faster with increasing depolarization (**Figure S3G**).

Finally, we compared AChR clustering, an integral step in NMJ development, in 2-week differentiated 2D and 3D muscle fiber cultures (**Figures 1C-G**). We observed significantly higher expression of the nAChR-β protein in 3D compared to 2D cultures at 2-weeks of fiber differentiation (**Figure 1C** and **Figures S2A and S2F**). Further, our analyses revealed a greater number of AChR clusters per muscle fiber established in 3D compared to 2D culture (**Figure 1E**). Indeed, we noted that at 2-weeks of culture, the majority of muscle fibers in 2D cultures lacked AChR clusters (**Figure 1A and 1E**). Interestingly, although average AChR cluster area was not significantly different (**Figure 1F**), we observed a high frequency of branched and perforated AChR clusters in our 3D muscle cultures, whereas oval shaped AChR clusters dominated on muscle fibers cultured in 2D conditions (**Figure 1D**). To quantify this observation, we assessed the lacunarity of AChR clusters formed on muscle fibers cultured in 2D and 3D conditions. Lacunarity is a measure of shape morphological heterogeneity and ‘gappiness’. Patterns with high lacunarity contain gaps or ‘lacunas’, whilst lower lacunarity implies pattern homogeneity or rotational invariance (Karperien et al., 2013; Smith et al., 1996). Lacunarity calculated from box counting validated our qualitative observations by indicating a significantly higher average lacunarity of AChR clusters formed in 3D cultures compared to 2D cultures (**Figure 1G**).

Overall our comparison of muscle fibers established in 2D and 3D formats suggests that a 3D culture method is ideal to support rapid contractile apparatus maturation and function, as well as AChR clustering and morphological maturation.

### 3D human neuromuscular co-cultures recapitulate early NMJ synaptogenesis

Since muscle fiber maturation is a prerequisite for NMJ development (Fox, 2009), we evaluated the hypothesis that the 3D skeletal muscle tissue platform would be well suited for human PSC-derived MN incorporation to model human NMJ synaptogenesis. We utilized MN clusters (Day 20) differentiated from WA09 human embryonic stem cell – derived OLIG2^+^ progenitor cells (**Figure S4A-B**) (Lippmann et al., 2015). Resulting MN clusters were enriched for cells expressing the HB9 and ISL1 transcription factors as well as the mature neurofilament marker SMI32 (**Figure S4C-D**). MN clusters were collected prior to muscle tissue preparation, mixed with the muscle progenitor cells in the hydrogel mix, and seeded together into the PDMS channels. The 3D skeletal muscle tissue media was optimized to support co-culture health by supplementation with brain derived and glial cell line derived neurotrophic factors (BDNF, GDNF) to support MN viability. Co-cultures examined after 10 days in differentiation media showed close contact between the MN clusters and the muscle tissue by phase-contrast microscopy (**Figure 2A**). Immunostaining co-cultures on the second week of culture for the motor neuron marker SMI-32, muscle fiber marker sarcomeric a-actinin, and a-bungarotoxin (to visualize AChRs) revealed that the co-cultures self-organized such that muscle progenitor cells fused to form multinucleated, aligned and striated muscle fibers and the MN clusters were positioned at the periphery of muscle bundles (**Figure 2B**). Importantly, the MNs were capable of regrowing neurites that were found in contact with a-bungarotoxin positive AChR clusters on muscle fibers (**Figure 2B-C**). In vivo studies by others found that postsynaptic AChR aggregation on muscle fibers is supported by agrin secretion from MN axon terminals (Gautam et al., 1996). We confirmed agrin expression in our PSC-derived MN cultures (**Figure S4E**). Furthermore, western blot analysis of neuromuscular co-cultures confirmed expression of MuSK (**Figure S5A**) and rapsyn (**Figure S5B**) proteins, two decisive synaptic proteins for mediating agrin-induced synaptogenesis (Glass and Yancopoulos, 1997). Consistently, we observed more and larger a-bungarotoxin positive AChR clusters in 3D neuromuscular co-cultures (**Figures 2D-F**), particularly at sites where MN neurites contacted muscle fibers (**Figure 2B**, yellow boxes). In addition, we observed a higher frequency of perforated and branched AChR clusters in our 3D co-cultures as evidenced by the higher lacunarity of AChR clusters formed in cocultures (**Figure 2G**) supporting a role for motor axon derived factors in post synaptic differentiation of the NMJ. As expected, by supplementing 3D human muscle tissue media with neural agrin (50 ng/mL) we phenocopied these co-culture results (**Figures 2H-J**). An evaluation of 2D neuromuscular co-cultures at the same time-point revealed a local alignment of the neurites and muscle fibers (**Figure S5C**, right panel), and a qualitative improvement in muscle fiber number and integrity (data not shown). However, only rare muscle fibers possessed clustered AChRs and we could not detect co-localization of the AChRs with SMI-32 stained neurites at this time point (**Figure S5C**).

**Figure 2.**
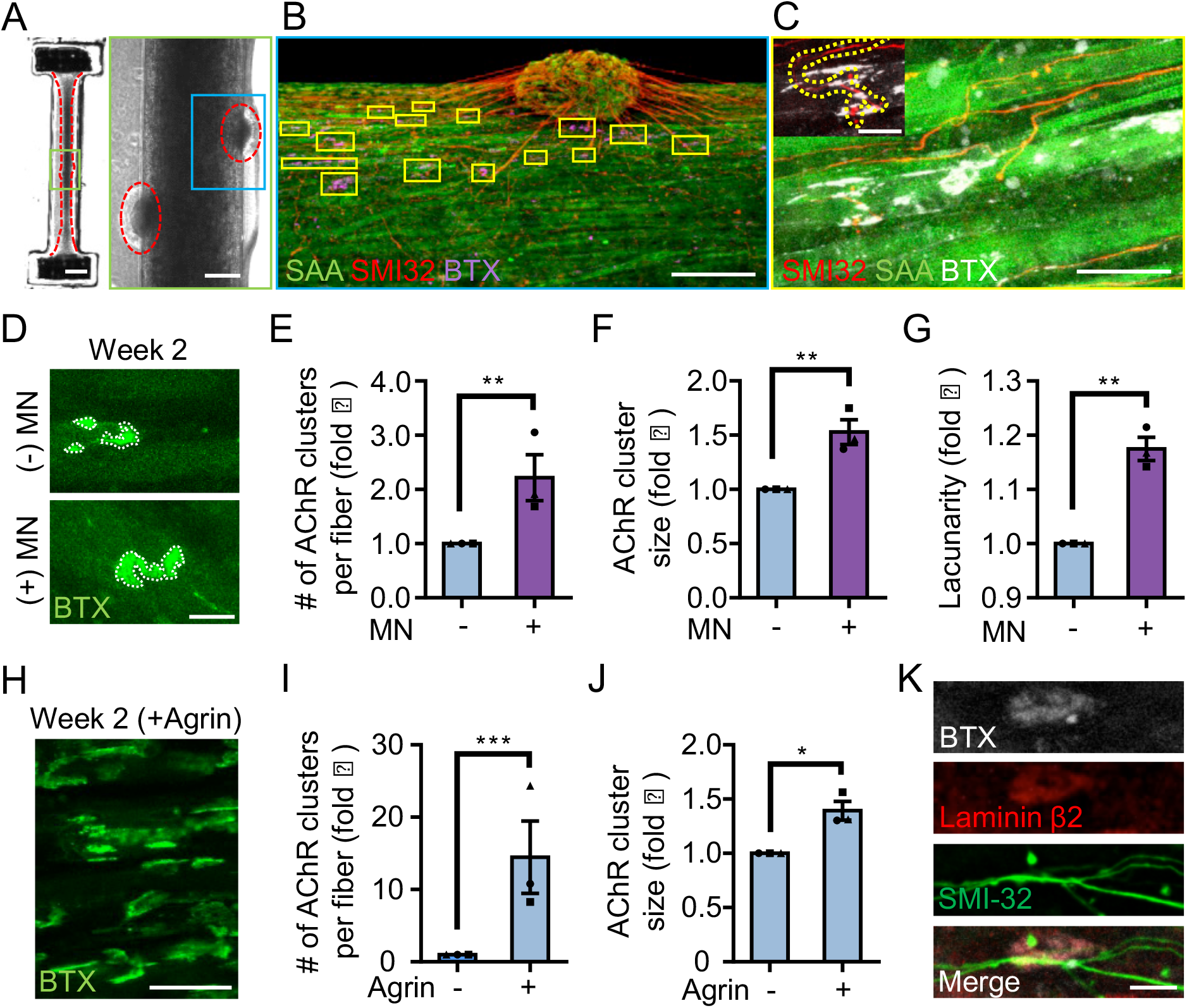
3D neuromuscular co-culture augments AChR clustering and maturation. **(A)** Stitched phase contrast image of a representative 3D skeletal muscle-motor neuron (MN) co-culture at two weeks of culture. Neuromuscular tissue outlined with red dashed line in left panel. Region outlined in green box is magnified in the image to the immediate right. Red dashed lines in right panel outline motor neuron clusters. Scale bars, 2 mm (left panel) and 200 μm (right panel). **(B)** Representative confocal image of a two-week old neuromuscular coculture immunostained for sarcomeric a-actinin (SAA; green), a-bungarotoxin (BTX; magenta), and neurofilament heavy SMI-32 (red). AChR clusters co-localized with neurites are outlined with yellow boxes. Scale bar, 200 μm. **(C)** Representative confocal image indicating colocalization of a SMI-32 (red) labeled neurite terminal and a BTX (white) labelled AChR cluster on a striated muscle fiber as seen by SAA (green) staining. Scale bar, 50 μm. **(D,H)** Representative confocal images of AChR clusters formed on muscle fibers cultured in 3D **(D)** with (+) or without (−) motor neurons (MN) or **(H)** supplemented with agrin and labeled with a-bungarotoxin after two weeks of culture. Scale bars, 25 μm (D) and 50 μm (H). AChR clusters are outlined with white dashed lines in **(D). (E-G, I-J)** Bar graphs indicating average **(E,I)** number of AChR clusters per fiber, **(F,J)** AChR cluster size, and **(G)** AChR cluster lacunarity in 3D cultures **(E-G)** with (+; purple) or without (-; blue) MN or **(I-J)** with or without agrin supplementation at week 2. In (E-G), values are normalized to 3D muscle cultures without MNs. In (I-J) values are normalized to untreated control. **(K)** Representative confocal image of a neuromuscular co-culture immunostained for laminin-β2 (red), bungarotoxin (BTX, white), or SMI-32 (green). Scale bars, 10 μm. For (E-G) and (I-J), n = minimum of 9 independent samples from 3 muscle patient donors. For agrin treated samples in (I-J), 6 samples from 3 muscle donors were analyzed. A minimum of 30 (E-G) or 6 (I-J) microscopic images per culture condition were analyzed. In (E-G) and (I-J) each symbol represents data from one muscle patient donor. Values in (E-G) and (I-J) are mean ± SEM. * p < 0.05, ** p < 0.01, and *** p < 0.001.

In further support of 3D muscle fiber synaptogenic maturation, the LAMB2 gene encoding for the laminin beta 2 chain was expressed by 3D human muscle tissues and neuromuscular co-cultures (**Figure S5D**), and the protein was found enriched at AChR clusters (**Figure 2K**). This is consistent with prior reports demonstrating laminin beta 2 concentrated at the neuromuscular junction synaptic cleft (Hunter et al., 1989) and the involvement of this tissue restricted basement membrane protein in NMJ maturation and maintenance (Noakes et al., 1995).

Our characterizations demonstrate that a 3D neuromuscular co-culture system, but not 2D co-cultures, recapitulates many aspects of early synaptogenesis that were first identified with in vivo studies.

### 3D human neuromuscular co-cultures are functionally innervated

We next sought to evaluate NMJ functionality in our neuromuscular co-cultures. With a combination of calcium handling analyses and electrophysiological recordings we report that 3D human neuromuscular co-cultures are functionally innervated in as early as two weeks. Using the fluorescent styryl dye FM 1-43 (Gaffield and Betz, 2007) and confocal microscopy we performed exocytosis assays on differentiated MNs (Day 20) and confirmed that human PSC-derived MNs exocytose in response to potassium chloride (KCl, 60 mM) and the excitatory neurotransmitter, L-glutamate (50 μM) stimuli (**Figures S6A-D** and **Movie S5**). The latter is particularly important, since the amino acid glutamate is a neurotransmitter that specifically stimulates MN cells but not muscle fibers (50 μM; **Movie S3**).

Next, we stimulated neuromuscular co-cultures that were generated using GCaMP6 transduced muscle progenitor cells with a 50 μM glutamate solution and observed calcium transients (**Figures 3A-B** and **Movies S6**) and synchronous tissue contractions (**Figure 3C** and **Movie S7**) in the muscle fibers in close proximity to the MN clusters in as early as 14 days of co-culture, indicating the formation of functional connectivity between MN endplates and muscle fibers. Stimulating the same tissue with ACh, following the glutamate stimulation, provided a rapid way to stimulate and visualize all muscle fibers in the tissue. Our analysis of this serial stimulation data revealed that many, but not all the fibers were functionally innervated (**Figure 3A-B** and **Movie S6**). As expected, direct stimulation of AChRs using ACh led to higher coculture contractile force generation as quantified by tissue movement (**Figure 3C**). To further validate that the presynaptic activation of motor neurons (i.e. glutamate stimulation) caused the observed changes in muscle fiber calcium transients and muscle fiber contractions, we studied the effect of BOTOX^®^ (BOT, presynaptic blocker) and d-tubocurarine (DTC, post synaptic blocker) treatments in our system. Our studies revealed a significant decrease in calcium transient activity and an absence of tissue contraction in response to glutamate stimulation if neuromuscular co-cultures were pre-treated with BOTOX^®^ or d-tubocurarine (**Figure 3D** and **Movie S7**).

**Figure 3.**
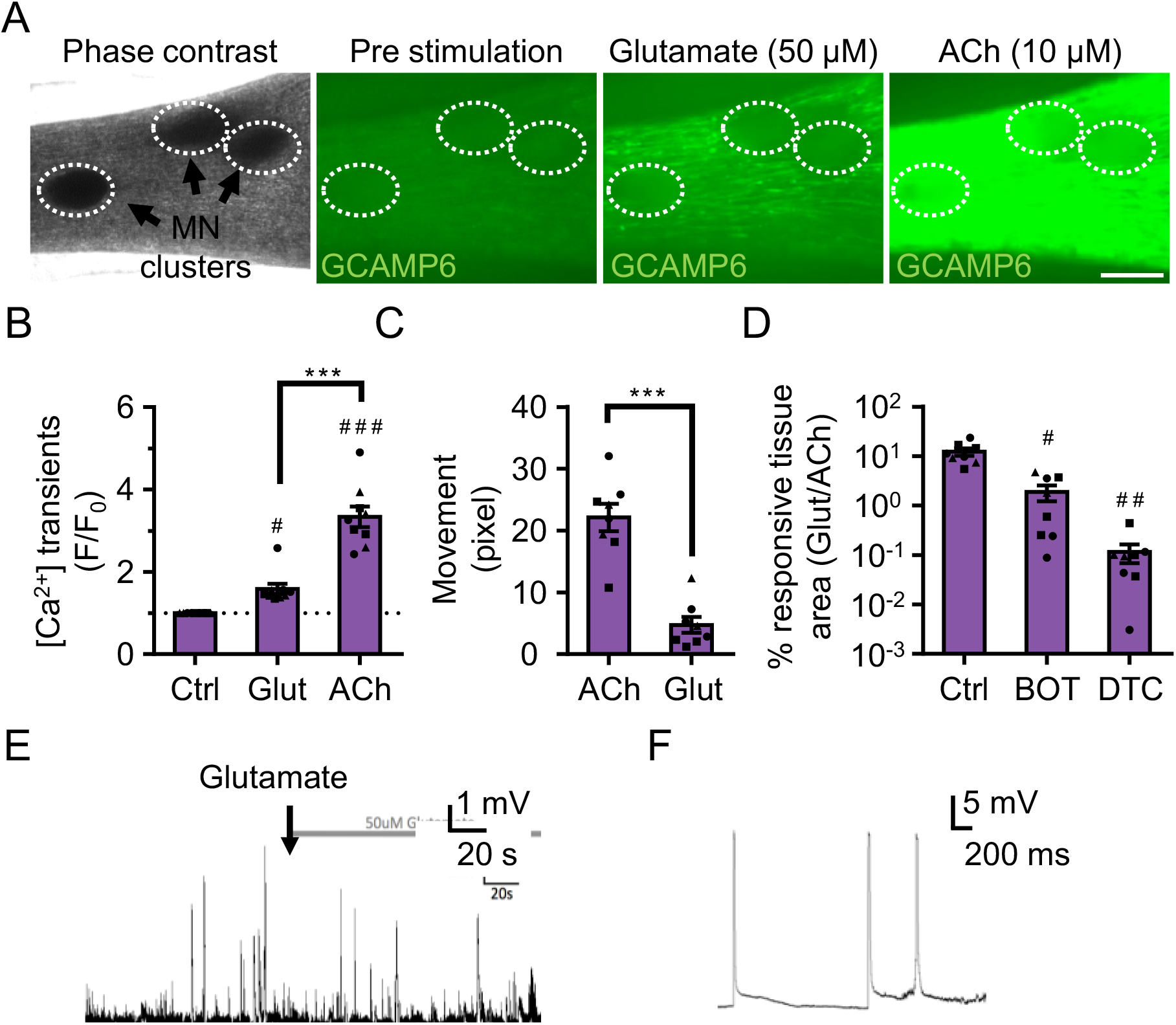
3D neuromuscular co-cultures are functionally innervated. **(A)** Phase contrast (far left panel) and GCaMP6 epifluorescence images (right panels) of a 3D neuromuscular co-culture after treatment with phosphate buffered saline (middle left panel), glutamate (middle right panel), or ACh (far right panel). Motor neuron clusters are outlined with white dashed lines. Scale bar, 250 μm. **(B)** Bar graph indicating quantification of fluorescence signal from neuromuscular co-cultures following glutamate (Glut) and Acetylcholine (ACh) stimulations relative to treatment with phosphate buffered saline (Ctrl). n = 9 neuromuscular coculture samples from 3 separate muscle patient donors. ^#^p < 0.05 and ^###^p < 0.001 compared with saline stimulation (Ctrl). **(C)** Quantification of neuromuscular co-culture tissue contraction in response to ACh (10 μM) and glutamate (50 μM). **(D)** Bar graph quantification of the percent tissue area occupied by glutamate (glut, 50 μM) responsive (GCaMP6^+^) fibers in saline (Ctrl), BOTOX^®^ (BOT, 1U), and d-tubocurarine (DTC, 25μM) treated 3D neuromuscular co-cultures. ^#^p < 0.05 and ^###^p < 0.001 compared with saline treated sample. In (C, D) n = 8 independent neuromuscular samples from 3 separate muscle patient donors. **(E)** Representative trace of a sharp microelectrode recording showing endogenous EPPs in a neuromuscular co-culture before and after addition of glutamate (arrow; 50 μM) to the bath media. **(F)** Representative action potential-like events recorded from a 3D neuromuscular co-culture following glutamate stimulation (right panel). In (B-D) each symbol represents data from one muscle patient donor. Values in (B-D) are mean ± SEM. ***p < 0.001.

In contrast, and as expected, we observed very few functional connections when evaluating 2D neuromuscular co-cultures matured for 2-weeks and then treated with glutamate (**Figure S6E** and **Movie S8**). Indeed, a prior report of 2D human neuromuscular co-cultures performed functional assays only after 60 days of culture (Steinbeck et al., 2016). To confirm that the muscle fibers possessed functional AChR channels, despite limited innervation at this time point, the 2D co-cultures were stimulated with ACh (100 μM). Calcium transients visualized by tracking GCaMP signal indicated the presence of ACh responsive muscle fibers in close proximity to the MN cluster (**Figure S6E** and **Movie S8**).

Next, to determine the maximum length of the functional connectivity between the MN cluster and the muscle fibers in 3D cultures, we generated neuromuscular tissues using GCaMP6 transduced muscle progenitor cells and a single MN cluster. On the second week of co-culture, calcium transients arising from the 3D neuromuscular tissues were recorded during glutamate (50 μM) stimulation. Analysis of pre- and post-stimulation movies indicated an average maximum functional connectivity length of 1042.7±104.5 μm at this time-point. As expected, the number of innervated fibers decreased as the distance from the MN cluster increased (**Figure S6F**).

Finally, we performed electrophysiological recording to directly address the functional properties of the neuromuscular junctions. Using current clamp, we observed spontaneous endogenous endplate potentials (EPPs) from single muscle fibers that were proximal to the MN cluster (**Figure 3E**), which were absent in muscle-alone cultures (**Figure S3E**). Upon glutamate stimulation, the frequency of EPPs was increased (**Figure S6G**), whereas the amplitude remained unchanged (**Figure S6H**). These results support the notion that the MNs were stimulated by glutamate to release neurotransmitter into the NMJ. Moreover, in these muscle fibers, we captured events that resembled action potentials in response to glutamate stimulation (**Figure 3F**). These events were characterized by a ~26 mV depolarization followed by a small plateau phase lasting ~8.5-14.5 milliseconds, but the absence of an afterhyperpolarization.

Together, these studies indicate that 3D neuromuscular co-cultures support efficient functional innervation that occurs faster than previously reported for 2D neuromuscular cocultures (Steinbeck et al., 2016).

### 3D human neuromuscular co-cultures to model adult NMJ development and disease

Next, given the high degree of innervation achieved in our neuromuscular co-cultures, we hypothesized that the 3D model might be capable of supporting the gamma (embryonic) to epsilon (adult)-subunit switch that was not observed in 2D human neuromuscular co-cultures (Steinbeck et al., 2016). Selective transcription of the AChR subunits occurs during different developmental stages (Martinou et al., 1991) and neural derived glycoprotein neuregulin-1 (NRG1), a motor neuron-derived factor, is thought to stimulate expression of the epsilon subunit of the AChR gene (CHNRE), which encodes an adult muscle AChR subunit (Ahn Jo et al., 1995; Falls et al., 1993). Using western blot experiments, we confirmed the expression of NRG1-β1 in our PSC-derived MNs (**Figure S7A**). Next, we quantified CHRNE expression in our 2D and 3D muscle-alone cultures and neuromuscular co-cultures. We observed a significant increase in the expression of the CHRNE gene in co-cultures compared to muscle-alone cultures, in both 2D and 3D after two weeks of culture (**Figures S7B**), suggesting involvement of MN-derived trophic factors in CHRNE gene expression. Given the limited innervation observed in 2D co-cultures at this time-point (**Figure S6E** and **Movie S8**), we speculate that NMJ-independent localized neurotrophic factor delivery contributes to CHRNE gene expression in muscle cells. To test whether the increase can be associated with NRG1-β1-mediated induction of the CHRNE gene, we supplemented our 2D and 3D muscle-alone cultures with recombinant NRG1-β1 (5 nM) and detected a significant increase in CHRNE expression in the supplemented muscle fiber cultures (**Figure S7B**). Treating 3D muscle-alone cultures with motor neuron-derived conditioned media did not induce epsilon subunit gene expression above untreated 3D muscle alone cultures (CHRNE 0.5% ± 0.4 % of GAPDH expression), suggesting that MN axon contact with muscle fibers may be necessary to locally deliver concentrated neurotrophic factors and modulate epsilon gene expression in muscle fibers.

We next evaluated AChR epsilon expression at the protein level and found that it was upregulated in 3D co-cultures, but not in 2D co-cultures (**Figures 4A-B**). The upregulation of AChR epsilon protein expression in 3D co-cultures was accompanied by a significant increase in AChR beta and no change in the AChR gamma subunit (**Figures 4A-B**), in support of studies concluding that gamma subunit transcription and translation does not appear to influence the onset or magnitude of epsilon expression (Witzemann et al., 1996; Yampolsky et al., 2008), and hinting that some embryonic AChRs may remain. MN-dependent changes in AChR subunit protein levels (beta, gamma, and epsilon) were not observed in 2D co-cultures (**Figure 4A**). These observations support the notion that AChR epsilon protein stability is influenced by the degree of muscle fiber and / or NMJ activity (Caroni et al., 1993; Missias et al., 1996; Witzemann et al., 2013; Xu and Salpeter, 1997; Yampolsky et al., 2008).

**Figure 4.**
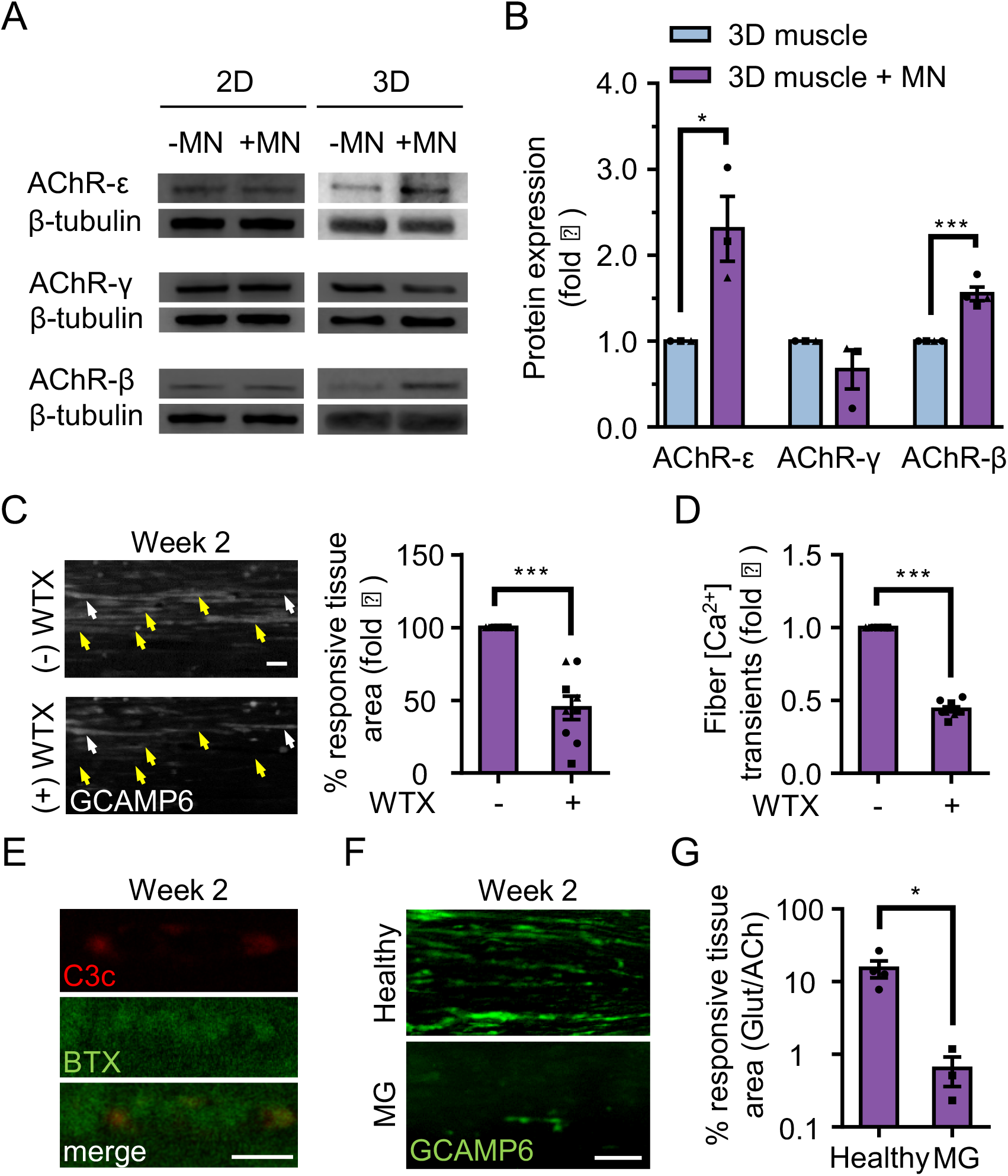
3D neuromuscular co-cultures enable disease modeling of adult NMJ in vitro. **(A)** Representative western blot images of nicotinic acetylcholine receptor subunit epsilon (nAChR-ε), gamma (nAChR-γ), and beta (nAChR-β) proteins in 2D and 3D muscle-alone (-MN) and neuromuscular co-cultures (+MN) at two weeks of culture. **(B)** Bar graph quantification of nACHR subunit ε, γ, and β protein expression in 3D muscle (blue) and 3D neuromuscular (purple) cultures. Values are normalized to 3D muscle cultures. **(C)** (left panel) Representative epifluorescence images of GCAMP6 signals in response to glutamate (glut) stimulation before (top panel) and after (bottom panel) 3D neuromuscular co-culture treatment with Waglerin 1 (WTX-1). Yellow arrowheads point out fibers with dampened GCAMP6 fluorescence signal following WTX-1 treatment. White arrowheads indicate fibers that did not dampen calcium handling after WTX-1 treatment. Scale bar, 50 μm. (right panel) Bar graph indicating the percentage of 3D neuromuscular co-culture tissue area occupied by glutamate responsive fibers (GCaMP6^+^) before (−) and after (+) WTX (1 μM) treatment. **(D)** Bar graph quantifying glutamate-induced GCAMP6 signals from individual fibers before (−) and after (+) WTX-1 treatment. In (CD), data is normalized to (−) WTX condition. For **(B-D)**, n = 9 independent muscle or neuromuscular samples from 3 muscle patient donors. A minimum of 50 fibers were analyzed for data presented in (D). **(E)** Representative confocal images of a 3D muscle culture co-treated with Myasthenia gravis (MG) patient IgG and human complement and then immunostained for human complement component C3c (red, top) and a-bungarotoxin (BTX, green, middle). Bottom panel is a merged image of the top and middle panels. Scale bars, 10 μm. **(F)** Representative epifluorescence images of GCaMP6 signals from a glutamate stimulated 3D neuromuscular co-culture following a 72-hour treatment with 300 nM of healthy (top panel) or MG (bottom panel) patient IgG and human complement. Scale bars, 100 μm. **(G)** Bar graph indicating the percent tissue area occupied by glutamate (glut, 50 μM) responsive (GCaMP6^+^) fibers in healthy and MG patient IgG treated 3D neuromuscular co-cultures. Data normalized to the total area of ACh responsive (GCaMP6^+^) tissue in each co-culture. n = 4 independent neuromuscular tissues treated with healthy IgG and 3 neuromuscular tissues each treated with serum IgG from one of three separate MG patient donors. In (B-D) and (G) each symbol represents data from one patient donor. Values in (B-D) and (G) are mean ± SEM. * p < 0.05 *** p < 0.001.

We then sought to determine if the 3D human neuromuscular co-culture system was suitable for modelling congenital myasthenic syndromes caused by mutations in CHRNE by blocking the AChR-epsilon subunit using Waglerin-1 (WTX); a peptide that selectively binds and blocks the epsilon subunit of the muscle AChR (McArdle et al., 1999). The AChR channel contains two binding sites for ACh, and one of those sites sits between the epsilon and a beta subunit in the adult AChR. Thus, if the epsilon subunit is functionally integrated into the AChR in neuromuscular co-cultures, then WTX treatment is expected to dampen calcium transients following glutamate stimulation by decreasing the statistical likelihood that the AChR channel will open (Jha and Auerbach, 2010; Ohno et al., 1996). In these experiments, 3D neuromuscular tissues were generated using GCaMP6 transduced muscle progenitor cells and each tissue was stimulated with glutamate twice: pre- and post WTX treatment (1 μM), with a 24-hour recovery time allocated between each stimulation. We recorded movies during glutamate stimulation and then quantified the maximal tissue area containing glutamate responsive fibers by analyzing the GCaMP6 fluorescence signal in the same tissue pre- and post-WTX treatment at defined regions of interests (**Figure 4C** and **Movie S9**). Consistently, we observed a 46.47 ± 15% (N = 3; p < 0.05) decrease in glutamate responsive tissue area following glutamate stimulation in WTX pre-treated neuromuscular tissues (**Figure 4C**). Similar results were obtained by analyzing calcium transients in individual fibers pre- and post WTX treatment in response to glutamate stimulation (**Figure 4D**; 55.8 ± 1.8% decrease). This analysis also revealed a subset of WTX-treatment refractory single fibers (**Figure 4C**, white arrowheads), indicating that not all AChRs in the 3D neuromuscular co-culture undergo the developmental switch by this time point. We performed similar experiments on 2D neuromuscular co-cultures (**Movie S8**), and 3D muscle-alone cultures (**Movie S10**), but did not observe calcium transient changes. Importantly, 3D neuromuscular tissue GCaMP signal was not dampened by serial glutamate stimulation (see methods) excluding the possibility that GCaMP dampening was the result of glutamate neurotoxicity.

Collectively, this data suggests that a 3D neuromuscular co-culture platform is ideal to rapidly and easily model and study diseases impacting the adult human NMJ.

### 3D human neuromuscular co-cultures to model myasthenia gravis

To demonstrate the tractability and robustness of the 3D neuromuscular co-culture system to study human disease, we treated co-culture tissues with IgG isolated from 3 patients afflicted with AChR-targeted myasthenia gravis (**Table S4**) to model autoimmune myasthenia gravis. Myasthenia gravis (MG) is an autoimmune disease manifesting as muscle weakness caused by the production of autoantibodies that alter, block, or destroy NMJ receptors required for signal transmission. IgG and complement deposit at the NMJ eliciting inflammation and subsequent destruction of AChRs on the postsynaptic NMJ membrane (Engel et al., 1977). Therefore, we treated our neuromuscular tissues with IgG (300 nM) isolated from healthy or MG patients together with human serum, which contains complement. Localized deposition of complement on BTX stained AChRs was confirmed by staining for the complement C3c protein one-day after co-treating muscle tissues with MG IgG and active human complement (**Figure 4E**). We recorded neuromuscular co-culture GCaMP signals arising from L-glutamate (50 μM) stimulation after a 3-day incubation with healthy or MG IgG (**Figure 4F**), to visualize NMJ activity. We then stimulated the co-cultures with ACh to quantify the total area occupied by muscle fibers. Our analysis revealed a clear decrease in the area of stimuli responsive muscle fibers (**Figure 4G** and **Movies S11-S12**) and a decline in overall tissue integrity (**Figures S8A-B** and **Movies S11-S12**) when tissues were treated with MG compared to healthy patient IgG.

This study demonstrates the simplicity of implementing the 3D neuromuscular co-culture system to the application of modeling and studying human NMJ disorders in culture.

## Discussion

Here we report the first method to co-culture 3D human skeletal muscle fiber tissues together with human PSC-derived MNs. We demonstrate that functional innervation is achieved in 3D, but not 2D neuromuscular co-cultures, within 2-weeks of culture. Indeed, we find that innervation in 3D neuromuscular co-cultures is ~4-fold faster and more efficient than a prior report of a 2D human neuromuscular co-culture system (Steinbeck et al., 2016), and we show that this simplifies and expedites studies of myasthenia gravis in a dish. With side-by-side comparisons of 2D and 3D muscle-alone and neuromuscular co-cultures, we confirmed that CHRNE transcription is supported by MN co-culture in 2D and 3D, and then show that the AChR epsilon subunit protein is only functionally integrated into the AChR in the context of 3D neuromuscular co-cultures. Therefore, this is the first report of a culture method to study the de novo AChR gamma to epsilon subunit developmental switch in culture and to model diseases of the adult human NMJ in a dish.

Our side-by-side comparison of human skeletal muscle fiber cultures in 2D and 3D indicates the structural and functional advantages of a 3D culture model over 2D systems. The value of 3D culture is reported in previous studies for other organs (Lancaster and Knoblich, 2014), and in this study we provide the first evidence that 3D culture conditions are indispensable for the maturation of multinucleated muscle fibers due to their capability to accommodate the inherent contractile nature of the muscle fibers in the long-term. This in turn leads to muscle fiber hypertrophy, improved calcium handling, and muscle fiber maturation as evidenced by expression of adult forms of MHC, and elaborated clustering of AChRs. This makes 3D neuromuscular cultures an ideal platform for studying NMJ synaptogenesis given the innately long process required for functional NMJ development to occur.

Consistently, electrophysiological recordings of single muscle fibers in these neuromuscular co-cultures detected endogenous and glutamate-stimulated EPPs, suggesting that MNs form functional neuromuscular junctions, similar to that observed in in vivo mammalian models. The properties of action potentials recorded in these cultures is consistent with the possibility that the 3D human neuromuscular co-cultures are not fully mature. This observation concurs with the relatively small muscle fibers, and indicates that additional chemical or physical cues are necessary to mature the tissues further. We saw a large action potential-like response in 1 out of the 7 fibers we assessed. This suggests that although the NMJs are functional, exhibit endogenous activity, and that motor neurons respond to glutamate application by increasing the basal rate of neurotransmitter release, most motor neurons are still in an immature state and do not trigger synchronous neurotransmitter release in response to glutamate application. This could be at the level of action potential generation in response to glutamate application, or converting action potentials to synchronous release at the presynapse. We anticipate increasing culture time, providing electrical stimulation, and / or adding trophic or synaptogenesis factors might improve the maturity of the neuromuscular co-culture and their connections.

Perhaps most strikingly, 3D neuromuscular cultures robustly produce functional adult AChR epsilon subunit, which is, to our knowledge, the first report of a system that supports the de novo gamma to epsilon AChR subunit switch in culture. In a proof-of-concept study, we demonstrate the application of our NMJ model to study adult NMJ activity by using a peptide that specifically blocks the epsilon subunit. Treatment with the peptide dampened glutamate-induced GCaMP6 calcium reporter activity in neuromuscular co-cultures demonstrating the utility of the system for adult NMJ studies.

Tissue culture affords the opportunity to deconstruct the complexity of a tissue system and to systematically rebuild complexity as a means to identify physical and chemical factors that influence biological processes. This method is particularly powerful in studies of the NMJ where decoupling nerve and muscle influences during development and in the adult, within the context of an animal model, is confounded by tissue death. Through an iterative comparison of 2D and 3D muscle alone and neuromuscular co-cultures, we found that CHRNE transcript expression is upregulated in both 2D and 3D neuromuscular co-cultures. Bathing 2D or 3D muscle fiber cultures with a high concentration of recombinant neuregulin-1 phenocopied the effect of MN co-culture on CHRNE transcript, but CHRNE transcript levels were not induced in 3D muscle-alone culture treated with conditioned media from MNs. Since our PSC-derived motor neurons express neuregulin-1 protein, but we do not observe appreciable NMJ activity in our 2D neuromuscular co-cultures, we speculate that if CHRNE transcript induction is reliant on NRG-1, then localized MN-mediated delivery of the protein may be necessary to achieve physiologically relevant concentrations of the protein, and that transmission via the NMJ is not required. Importantly, epsilon protein levels further increased and its function was detected (WTX-responsivity) only in the context of 3D neuromuscular co-cultures. Our culture data indicates that the epsilon subunit of the AChR is subjected to post-transcriptional modifications and/or intracellular trafficking events that are only supported in the context of 3D neuromuscular co-culture. Indeed, our observations that muscle fibers established in 3D culture are more mature (**Figure 1 and Figures S1-S2**) and that 3D neuromuscular co-cultures exhibit spontaneous endogenous endplate potentials (**Figure 3 and Movie S6**) fit well with studies linking muscle fiber maturation state and activity to AChR subunit conversion and stability (Caroni et al., 1993; Missias et al., 1996; Witzemann et al., 2013; Xu and Salpeter, 1997; Yampolsky et al., 2008). With delivery of this new methodology supporting de novo adult NMJ development, it is now possible to delve deeper into the mechanisms regulating epsilon activity in normal development and in disease states.

In summary, our method to model the adult human NMJ in a dish provides a versatile approach to study skeletal muscle and NMJ development, but more importantly, provides the first reproducible method to study adult, rather than embryonic, NMJ activity in as early as two weeks of co-culture time. Our calcium reporter neuromuscular tissues can easily be integrated with other optogenetic methods (Steinbeck et al., 2016) to further elucidate synaptic transmission mechanisms in adult NMJ, such as adult AChR conductance. Furthermore, neuromuscular co-cultures may be integrated with other neuron populations such as upper MNs and / or myelinating Schwann cells to support studies aimed at a better understanding of signal transmission in the central nervous system. The method now enables modelling diseases that target the adult NMJ (e.g. congenital myasthenia gravis, Duchenne muscular dystrophy (Xu and Salpeter, 1997) as well as MN dependent disorders of the NMJ (e.g. spinal muscular atrophy and amyotrophic lateral sclerosis). Finally, the 3D human NMJ system can serve as an improved pre-clinical pharmacological testing platform to evaluate drugs designed for NMJ disorders or to support personalized medicine applications.

## Supporting information

## Experimental Procedures

### Human primary myoblast derivation and propagation

Human skeletal muscle tissues removed in the course of scheduled surgical procedures and designated for disposal were utilized in this study in accordance with St. Michael’s Hospital research ethics board and University of Toronto administrative ethics review approval. Small skeletal muscle samples (~1 cm^3^) were obtained from the multifudus muscle of patients undergoing lumbar spine surgery. Primary myoblast and fibroblast-like cell lines were established and maintained as previously described (Blau and Webster, 1981). Briefly, human skeletal muscle samples were minced and then dissociated into a single cell slurry with clostridium histolyticum collagenase (Sigma, 630 U/mL) and dispase (Roche, 0.03 U/mL) in Dulbecco’s Modified Eagle’s medium (DMEM; Gibco). The cell suspension was passed multiple times through a 20 G needle to facilitate the release of the mononucleated cell population and subsequently depleted of red blood cells with a brief incubation in red blood cell lysis buffer (**Table S2**). The resulting cell suspension containing a mixed population of myoblasts and fibroblast-like cells was plated in a collagen-coated tissue culture dish containing myoblast growth medium: F-10 media (Life Technologies), 20% fetal bovine serum (Gibco), 5 ng/mL basic fibroblast growth factor (bFGF; ImmunoTools) and 1% penicillin-streptomycin (Life Technologies). After one passage, the cell culture mixture was stained with an antibody recognizing the neural cell adhesion molecule (NCAM/CD56; BD Pharmingen; **Table S1**), and the myogenic progenitor (CD56^+^) and fibroblast-like cell (CD56^−^) populations were separated and purified using fluorescence-activated cell sorting (FACS) and maintained on collagen coated dishes in growth medium. Subsequent experiments utilized low passage cultures (P4—P9).

### Human primary myoblast two-dimensional culture

Primary human myoblasts were mixed with primary human muscle fibroblast-like cells at the following ratios: CD56^+^ (95%) and CD56^—^ (5%). For Geltrex™ culture dish coating, 1 mg of Geltrex™ was resuspended in 12 mL of ice-cold DMEM and 1 mL was transferred to each well of a 12 well plate. Plates were incubated at 37 °C overnight. DMEM was aspirated the next day just prior to cell culture. 3×10^6^ cells resuspended in bFGF-free myoblast growth media (**Table S2**) were plated into each Geltrex™ (Life Technologies) coated well. The growth media was exchanged 2 days later with myoblast differentiation medium (**Table S2**). Half of the culture media was exchanged every other day thereafter. In some experiments (Supplemental Figures 2G-H), fibrinogen was supplemented into the differentiation media at 10 μg/mL to control for the effect of fibrinogen receptor ligation on two-dimensional (2D) muscle fiber differentiation.

### PDMS mold fabrication for 3D human muscle tissue culture

Standard 12-well culture plates were coated with 500 μL of liquid PDMS (184 Silicone Elastomer Kit, 10 parts elastomer to 1 part curing). After curing at 50 °C for at least 3 hours, another 750 μl of liquid PDMS was added to each well and a laser cut, dumbbell shaped piece of acrylic (middle channel dimensions = 14 mm by 2.75 mm; side chamber dimensions = 5.7 mm by 2.5 mm) was submerged in the liquid PDMS. Plates were then placed within a vacuum chamber for a minimum of 10 minutes to remove bubbles from the liquid PDMS. The PDMS was cured by incubating the plates in a 50 °C oven for 3 hours. Acrylic pieces were then removed from the PDMS, leaving a dumbbell-shaped depression in the PDMS, and two pieces of Velcro™ fabric were affixed at each end of the channel using liquid PDMS as glue (**Figure S1D**). Each well was sterilized with 70% ethanol at room temperature in a tissue culture hood for at least 30 minutes. At this point, plates were parafilm sealed, and stored at room temperature. Prior to use, PDMS mold wells were incubated with a 5% pluronic acid (Sigma) solution in ddH2O for 12 hours at 4 °C. Pluronic acid solution was aspirated and molds were rinsed with a PBS solution before seeding human muscle tissues.

### Human myoblast three-dimensional culture

Three-dimensional (3D) human skeletal muscle tissues were generated in culture as previously described (Madden et al., 2015) with the following modification: FACS-purified CD56^+^ myoblasts (95%) and CD56^−^ fibroblast-like cells (5%) were incorporated into tissues. Briefly, cells at these defined ratios were resuspended in the hydrogel mixture (**Table S2**) in the absence of thrombin. Thrombin (Sigma) was added at 0.2 unit per mg of fibrinogen just prior to evenly seeding the cell/hydrogel suspension in the long channel of the dumbbell-shaped molds. Tissues were then incubated for 5 minutes at 37 °C to expedite fibrin polymerization. Myoblast growth media (**Table S2**) lacking bFGF, but containing 1.5 mg/mL 6-aminocaproic acid (ACA; Sigma), was added. 2 days later the growth media was exchanged to myoblast differentiation medium (**Table S2**) containing 2 mg/mL ACA. Half of the culture media was exchanged every other day thereafter. In agrin treatment experiments, recombinant rat agrin (R&D Systems) was supplemented in the culture media at 50 ng/ml. In experiments using neuregulin1-β1 treatment, recombinant human neuregulin1-β1 (R&D Systems) was supplemented in the culture media at 5 nM. In both cases (agrin, neuregulin), treatment began when tissues were switched to differentiation medium and recombinant proteins were added to exchange media at 2-fold concentration.

### hESC differentiation to post-mitotic motor neurons

Motor neurons were specified from WA09 hESCs (passage 25-45; WiCell) as previously described (Lippmann et al., 2014, 2015). Briefly, hESCs were maintained on Matrigel (BD Biosciences) in E8 medium with insulin added at a concentration of 2 mg / L. For differentiation, hESCs were dissociated with accutase (Life Technologies) and reseeded at 1×10^5^ cells / cm^2^ in E8 medium containing 10 μM ROCK inhibitor (Y27632; R&D Systems) in 6-well polystyrene tissue culture plates coated with 100 μg / mL poly-L-ornithine (PLO; Sigma) and 8 μg / well VTN-NC (gift from Dr. James Thomson). hESC were differentiated to OLIG2^+^ progenitors in E6 medium containing the same insulin concentration as in E8 medium as previously described (Lippmann et al., 2015). For differentiation of the OLIG2^+^ progenitors to motor neurons, cells were sub-cultured by en bloc passage, reseeded at a 1:200 ratio in Geltrex^™^-coated 6-well plates, and differentiated for 14 days in E6 medium containing 1 μM retinoic acid (Sigma), 100 nM purmorphamine (Tocris), and 100 ng/mL sonic hedgehog (R&D systems). To push neuronal maturation before myoblast co-culture, 5 μM DAPT (Tocris) was added from days 8-14.

For a subset of experiments (**Figures 3C-D** and **Movie S7**), OLIG2^+^ progenitor cells were specified following a previously described method (Du et al., 2015) using GFP-expressing iPSCs (Nazareth et al., 2013). Next, OLIG2^+^ progenitors were differentiated to post mitotic motor neurons following the same differentiation protocol mentioned above.

### Two- and three-dimensional neuromuscular co-culture

24 hours after seeding myogenic progenitor cells for culture in 2D (as described above), 5 ESC-derived motor neuron clusters were detached and transferred to the muscle cell culture plates using a 1 ml pipette tip in myoblast media lacking bFGF, but now containing 10 ng / ml brain derived neurotrophic factor (BDNF) and 10 ng / ml glial cell line derived neurotrophic factor (GDNF). Mid-sized clusters (150 to 300 μm in diameter) were visually identified and selected for transfer. 24 hours later the media was removed and replaced with myogenic differentiation media (**Table S2**) supplemented with 10 ng / mL BDNF and 10 ng / mL GDNF. Half of the culture media was exchanged every other day thereafter and included both neurotrophic factors at 2-fold concentration.

For 3D neuromuscular tissues, 3D skeletal muscle tissue cell / hydrogel suspension was prepared as described above. Motor neuron clusters were transferred manually to the cell / hydrogel suspension at a ratio of 5 clusters per tissue. Thrombin was added and tissues were seeded into dumbbell-shaped molds as described above. Myoblast growth media lacking bFGF, but containing 2 mg / mL 6-aminocaproic acid (ACA; Sigma), 10 ng / mL BDNF, and GDNF was added to tissues. Two days later the culture media was exchanged to myogenic differentiation media (**Table S2**) and supplemented with 10 ng / mL BDNF and 10 ng / mL GDNF. Half of the culture media was exchanged every other day thereafter and included both neurotrophic factors at 2-fold concentration. 2D and 3D muscle-alone cultures serving as neuromuscular co-culture controls were also supplemented with BDNF and GDNF. Co-cultures were analyzed at time points indicated in the figures and legends.

For a subset of experiments (**Figures 4E-G, Figure S5C, Figure S6E, Figure S8A-B**, and **Movies S11-S12**), immortalized myogenic progenitor cells (AB1167, from fascia lata muscle of a healthy 20-year old male), were employed. Briefly, human-derived skeletal muscle cell (hSMC) lines used in this work were derived from healthy subjects and were then immortalized by transduction with human telomerase-expressing and cyclin-dependent kinase 4-expressing vectors, as previously described (Mamchaoui et al., 2011). For these experiments we transduced the immortalized human myogenic progenitor cells with lentiviral particles to express GCaMP6, a fluorescent calcium indicator (AddGene plasmid #65042; Trono Lab packaging and envelope plasmids, Addgene plasmid #12260 and 12259), as described on the Broad Institute website (https://portals.broadinstitute.org/gpp/public/resources/protocols). The cell population was then sorted for GFP expression to enrich transduced cells and were then further expanded in myoblast growth media (**Table S2**). Methods to produce 3D neuromuscular tissues using immortalized myogenic progenitor cells were exactly as those described above for primary human muscle progenitors with the exception that fibroblast-like cells were excluded.

### Immunostaining and fluorescence microscopy

2D cultures and 3D tissue whole mounts were fixed in 4% PFA for 10 minutes and then washed with phosphate buffered saline (PBS). Following fixation, samples were incubated in blocking solution (**Table S2**) for at least 1 hour. Samples were incubated in primary antibody solutions diluted in blocking solution (**Table S1**) overnight at 4 °C. After several washes in blocking solution, samples were incubated with appropriate secondary antibodies diluted in the blocking solution for 30 minutes at room temperature. Hoechst 33342 or DRAQ5 (ThermoFisher) were used to counterstain cell nuclei. Confocal images were acquired with Fluoview-10 software using an Olympus IX83 inverted microscope. Epifluorescence images were acquired with CellSense™ software using an Olympus IX83 microscope equipped with a Olympus DP80 dual CCD color and monochrome camera. Images were analyzed and prepared for publication using NIH ImageJ software.

### Myofiber size analysis

Myofiber size was measured by assessing 40X magnification confocal images of 2D and 3D cultures immunostained for sarcomeric α-actinin. 2D muscle culture images and flattened z-stack images of 3D muscle tissues were analyzed to quantify the diameter of each muscle fiber using the NIH ImageJ.

### Western blotting

3D tissues were collected at the indicated time points and flash frozen in liquid nitrogen, while 2D cell cultures were directly lysed in RIPA buffer (see below) and flash frozen. Processed 2D and 3D samples were stored at -80 °C until all desired time points were collected. Tissues and 2D samples were lysed in RIPA buffer (ThermoFisher) containing protease inhibitors, and then lysates were analyzed for total protein concentration using the BCA protein assay kit (ThermoFisher). 15 μg of protein was analyzed on an 8 % SDS PAGE gel. Western blot was performed using a Bio-Rad Power Pac 1000 and Trans-Blot Turbo Transfer System to transfer the proteins from the polyacrylamide gel to a nitrocellulose membrane. Primary antibodies (**Table S1**) were incubated with membranes overnight at 4 °C in milk-based blocking solution (**Table S2**). Membranes were washed 3 × 30 minutes with rocking in a Tris-buffered saline with Tween (TBST; **Table S2**) and then transferred into blocking solution containing horseradish peroxidase conjugated anti-rabbit and anti-mouse secondary antibodies (Cell Signaling; 1:5000). Chemoluminescence was performed using ECL substrate (ThermoFisher) with a MicroChemi 4.2 chemiluminescence imaging system (DNR Bio-Imaging Systems). Images were analyzed using the NIH ImageJ.

### AChR cluster analysis

α-bungarotoxin staining was performed to visualize and quantify the number, size, and morphology of AChR clusters in 2D cell and 3D tissue cultures. Briefly, fixed tissues were incubated with 5 nM of Alexa Fluor 647 conjugated α-bungarotoxin for 30 minutes to label AChRs. Samples were then washed with PBS and 40X images, all 0.1 mm^2^ in area, were captured at a minimum of 6 random locations per sample. AChR cluster outlines in each 40X image were generated using the ImageJ particle analyzer. Clusters smaller than 5 μm^2^ were excluded from analysis. AChR cluster outline drawings were binarized to facilitate downstream analysis. To assess cluster number, AChR cluster were quantified for each image and was then normalized to the number of sarcomeric a-actinin+ fiber units present in the quantified image. To assess cluster area, the area of each individual AChR cluster was measured and averaged for each experiment using NIH ImageJ software. Fractal analysis was performed on α-bungarotoxin stained sample images to quantify AChR cluster morphological differences across the different culture conditions. After binarization of the AChR cluster outline drawings, lacunarity, a measure of gappiness and heterogeneity in a shape, was measured for each AChR cluster using the NIH ImageJ FracLac plug-in. The Sub Sample and Particle Analyzer method was used with FracLac along with default settings and 4 grid locations. Lacunarity was measured for each AChR cluster within an experiment and then averaged.

### Electrical stimulation

To ensure accurate and reproducible conditions for electrical stimulation, a custom-made stimulation chamber was produced using a 35 mm petri-dish, 2 carbon rods, and platinum wires. Before each use the stimulation chamber was sterilized using 70% ethanol. At day 14 of differentiation, an individual tissue was transferred to the chamber and covered in differentiation medium. Platinum wires were hooked up to a commercial function generator (Rigol DG1022U). A Rigol DS1102E digital oscilloscope was used to confirm the frequency and amplitude of signals before connecting the pulse generator to the platinum wires. 3D tissues were stimulated using square pulses with 20% duty cycle, 5V amplitude (field strength of 1.67 V/cm), and the reported frequencies.

### Calcium transient analysis

CD56^+^ sorted human myogenic progenitor cells were transduced with a lentiviral vector encoding the fluorescent calcium indicator GCaMP6 driven by the muscle specific gene MHCK7 (AddGene plasmid #65042). Cells were then sorted to purify the infected cells based on GFP expression. Human skeletal muscle progenitor cultures expressing GCaMP6 were imaged using an Olympus IX83 microscope equipped with modules to control the temperature and CO_2_ concentration. Movies were recorded at 4X magnification at 12 frames per second under physiological condition (37 °C and 5% CO_2_) in differentiation media using an Olympus DP80 dual CCD color and monochrome camera and CellSense™ software. Acetylcholine (BIOBASIC) was reconstituted to produce a 100 mM stock solution in PBS and was diluted to the final working concentration (as specified in the text) by addition directly into the culture chamber.

In glutamate stimulation studies, L-glutamate (Abcam) was first reconstituted to 100 mM in 1equal NaOH and then further diluted in HBSS (Gibco) / DMEM to produce a 100x stock solution (5 mM). For AChR epsilon subunit blocking studies, Waglerin-1 (Smartox Biotechnology) was prepared as a 100x stock solution in PBS and was added to cultures at a 1 μM working concentration 10 minutes prior to stimulation.

To assess the effect of Waglerin-1 on glutamate-stimulated calcium transients, a movie was recorded for each tissue before and after glutamate stimulation. Video segments, equal in length, representing pre- and post-glutamate GcAMP6 signals were each projected into a 2D image. The 2D projected images were then subtracted to eliminate spontaneously active fibers from our analysis. Background from different imaging sessions were normalized. In Figure 4C, GCaMP6 signals were analyzed to quantify the area of glutamate responsive tissue at the same ROIs before (−) and after (+) Waglerin-1 treatment and presented as a fold-change. In Figure 4D, we identified all individual fibers that demonstrated GcAMP6 signal dampening in response to Waglerin treatment and then quantified signal in those fibers before (−) and then after (+) treatment and presented the data as a fold-change.

To assess the effect of BOTOX^®^ (Allergan, Irvine, CA) and d-tubocurarine (Sigma) treatments on glutamate-stimulated calcium transients, co-cultures were treated with BOTOX^®^ (1U) and d-tubocurarine (25 μM) for at least 10 minutes before glutamate stimulation in their culture media. BOTOX^®^ was prepared at 100 U/ml in PBS and d-tubocurarine was reconstituted at 2.5 mM in DMEM.

### Tissue contraction quantification

To assess the contraction of neuromuscular tissues following glutamate and acetylcholine stimulations, co-cultures were stimulated under the indicated experimental conditions and movies were recorded at 4X magnification at 12 frames per second under physiological conditions (37 °C and 5% CO_2_). To quantify neuromuscular tissue contraction, movies were assembled into stacks using ImageJ software and 3 regions of interest were traced within each stack. Maximum movement distance for each trace was determined and averaged for each sample. Data are presented as movement (distance) in pixels.

### Length of functional connectivity between MN cluster and muscle fibers

In this studies, neuromuscular co-cultures were generated using GCaMP6 transduced human myogenic progenitor cells and a single motor neuron cluster. At week two of co-culture, tissues were stimulated by a 50 μM L-glutamate solution diluted in DMEM. Movies were captured at a frequency of 12 frames per second for at least 15 seconds before and after stimulation. Movies were processed exactly as described above (calcium transient analysis) to eliminate the spontaneously active fibers from the analysis. The location of the motor neuron cluster was identified from a bright field image, which was used to outline the structure with a circle in the epifluroscent images. Using NIH ImageJ software, concentric circles, each 100 μm larger in radius than the prior, were outlined around the motor neuron cluster until the circles encompassed all the glutamate responsive fibers (i.e. GcAMP6^+^) on the subtracted image. The number of active fibers in each circle were then quantified to determine the number of fibers in each concentric circle ‘bin’. Binned data from 3 independent experiments were then reported on a histogram to report the average number of glutamate responsive fibers as it relates to the distance (i.e. concentric circle bin) from the motor neuron cluster.

### Electrophysiological recordings

Individual muscle fibers were impaled with 30-40 MΩ sharp electrodes pulled from borosilicate glass (World Precision Instruments), filled with 3M KCl. Membrane potential was recorded in the current clamp configuration using a Digidata 1440A and MultiClamp 700 A amplifier (Axon Instruments, Molecular Devices). Data were digitized at 10 kHz and filtered at 2.6 kHz. Data were quantified using MiniAnalysis (Synaptosoft). Each fiber was allowed to recover for a few minutes, to allow its resting membrane potential to stabilize before recordings were performed. For electrophysiological recordings following optogenetic stimulation, 3D muscle tissues were generated using human skeletal muscle progenitors transduced with a lentiviral vector encoding humanized ChR2 with H134R mutation fused to EYFP and driven by EF1a (AddGene plasmid #20942). Cells were sorted to purify the infected cells based on the EYFP signal. Optogenetic stimulation was performed using blue LED (KSL-70, RAPP OptoElectronic) with a wavelength of 470 nm, and controlled by the Axon amplifier software. In glutamate stimulation experiments, glutamate was pipetted by hand into the edge of the bath and allowed to diffuse to the tissue. For all recordings, the bath solution was standard DMEM (Gibco).

### Myasthenia gravis disease modeling

Serum from 3 patients diagnosed with Anti-AChR MG (**Table S4**) was collected and IgG fractions were purified using a Protein A IgG purification kit (Thermofisher) based on the manufacturers instruction. Purified IgG was reconstituted in PBS and IgG content was quantified using a NanoDrop 1000 spectrophotometer (ThermoFisher). On Day 11 of neuromuscular coculture, IgG was added to the differentiation media at 300 nM final concentration and 2% human serum (Sigma) was supplemented in the differentiation medium rather than horse serum. IgG from healthy human serum (Sigma) was used in ‘healthy’ control experiments. In these experiments, IgG was added once and the media was not exchanged thereafter. After 3 days of treatment, the Day 14 co-cultures were stimulated with glutamate followed by an acetylcholine stimulation and calcium transients were captured by imaging the GCaMP6 signals using an Olympus IX83 microscope. Video segments, equal in length, representing GCaMP6 signals from glutamate and ACh serially stimulated tissues were each projected into a 2D image. For Figure S8B, GCaMP6 signals were analyzed to quantify the area of ACh responsive tissue in equal sized ROI for healthy compared to MG IgG treated tissues, and presented as fold change. For Figure 4G, healthy and MG IgG treated tissues were analyzed to quantify the area of GCaMP6 signal at the same ROIs after glutamate (glut) and then acetylcholine (ACh) stimulation and the ratio of glutamate-to ACh-induced GCaMP6 signals was reported.

In a subset of experiments, we performed serial glutamate stimulation experiments in which we stimulated the same neuromuscular co-culture on Day 11 and then Day 14 of culture with a 50 μM L-glutamate solution and measured the area of GCaMP6^+^ tissue as described above. Our analysis indicated that the area of glutamate responsive tissue increased from ~2 % at Day 11 to more than 13 % on Day 14, arguing against glutamate-induced cytotoxic effects arising from our experimental methods.

### FM 1-43 labelling and imaging

Motor neuron clusters were separated from undifferentiated single cells using Accutase (ThermoFisher) and transferred to a Geltrex™ coated 6-well plate on Day 14 of differentiation. Clusters were then cultured for an additional week in E6 media supplemented with 10 ng/mL BDNF and 10 ng/mL GDNF, to permit the regrowth of the neurites, and were then labelled with the FM 1-43 styryl dye (Molecular Probes) following the manufacturer’s instructions. Briefly, MN clusters were stimulated using high potassium solution (60 mM) and incubated with FM 1-43 (2 μM) in HBSS (+Mg^2+^ and +Ca^2+^) for 20 minutes to enable dye loading. Clusters were washed with HBSS for at least one hour at room temperature before imaging. Samples were imaged using an IX83 Olympus confocal microscope with FV-10 software at physiological conditions (37 °C and 5% CO_2_). MN clusters were then stimulated with either high potassium solution (60 mM in PBS), L-glutamate (50 μM in HBSS), or control solutions (HBSS or PBS) while acquiring time-lapse movie sequences. Movies were analysed for fluorescence intensity before and after stimulation at each indicated time-point using NIH ImageJ software.

### Gene expression analysis

Total RNA was extracted from 3 technical replicate muscle tissues or neuromuscular co-culture for each of 3 biological replicate experiments using the PureLink RNA Micro Kit according to the manufacturer’s protocol (ThermoFisher). cDNA was reverse transcribed from 400 ng of RNA using the High-Capacity cDNA Reverse Transcription kit (Applied Biosystems). For quantitative real-time PCR (qRT-PCR), CHRNE and CHRNG primers were acquired from Bio-Rad and reactions were run according to manufacturer’s protocol on the Roche LightCycler 480 (Roche) using LightCycler 480 SYBR Green I Master (Roche). All results were normalized to the housekeeping gene glyceraldehyde 3-phosphate dehydrogenase (GAPDH). Gene expression is reported in % of GAPDH expression ± SEM. To assess agrin gene expression in differentiated MNs, cDNA samples were prepared from 3 consecutive MN differentiations. Genes were amplified using Arktik thermal cycler according to the manufacturer’s protocol (ThermoFisher). PCR amplification products were analyzed on a 2% agarose gel with SYBR safe DNA gel stain (Invitrogen). GAPDH gene expression served as the loading control. All oligo sequences are summarized in **Table S3**.

### Statistical Analysis

Each study in this manuscript was performed using three primary myoblast lines derived from three separate muscle patient donors (N = 3 biological replicates). Each experiment within a study was set-up with cells from a separate muscle donor and included at least n = 3 technical replicates. Exceptions include Figure 2I-J, Figure 4G, Figure S2G, Figure S3C, and Figure S8B. In Figures 2I-J, two muscle tissue cultures (technical replicates) were treated with Agrin for each biological replicate (6 samples in total). In Figure 4G and Figure S8B, IgG purified from sera collected from 3 MG patients was tested (N = 3 biological replicates), and compared to IgG purified from a single healthy donor and tested on 3D neuromuscular tissues engineered using a single immortalized cell line. In Figure S2G, 7 technical replicates from 3 muscle patient donors was analyzed at the 1-week culture time point. In Figure S3C, one muscle sample was analyzed for each muscle patient donor (N = 3 muscle patient donors). For all other neuromuscular co-culture studies, each primary myoblast line was co-cultured with MNs established from separate human pluripotent stem cell derivations.

Statistical analysis was performed on data obtained from technical replicates using GraphPad Prism 6.0 software. Statistical differences between experimental groups were determined in most studies by unpaired t-test. Exceptions to this are as follows: Two-way ANOVA followed by Tukey’s and Sidak’s multiple comparisons were performed in Figure 1B and Figure S7B. One-way ANOVA followed by Tukey’s multiple comparisons was performed in Figure 3B, Figure 3D, Figure S2G, Figure S3C, and Figure S6F. Results are presented as mean ± SEM. p < 0.05 was considered significant for all statistical tests. Absence of a significance symbol (^*^, ^#^, ^$^) indicates no significant differences.

## Acknowledgements

We would like to extend our thanks to Saint Michael’s Hospital and Telethon Network of Genetic BioBanks (GTB12001D) patients who agreed to contribute sera samples to our study. We would also like to thank the following sources for funding this study: NSERC CREATE TOeP, Toronto Musculoskeletal Centre, Ontario Graduate Scholarship, and Krembil Foundation to MAB; Postdoctoral NRSA from the NIH (1F32NS083291-01A1) to ESL; TD Bank Health Research Fellowships at the LTRI (to BM); the Myobank-AFM and the platform for immortalization of human cells at the Myology Institute (Paris, France; to AB); the Canadian Institute of Health Research Foundation Scheme 154274 to (MZ) and Natural Sciences and Engineering Research Council to (RGPIN-2017-06738 to MZ, BAS, and RGPIN 435724-13 to PMG); an Innovation in Regulatory Science Award for the Burroughs Wellcome Fund (RSA), NIH grant R21NS082618 (RSA), and this publication was developed, in part, with STAR center grant 83573701 to RSA from the U.S. Environmental Protection Agency (EPA) but has not been formally reviewed by the EPA. The views expressed in this document are solely those of RSA and colleagues and do not necessarily reflect those of the Agency, and the EPA does not endorse any products or commercial services mentioned in this publication; and Toronto Western Arthritis Program, Ontario Research Fund (31390), Canada Research Chair Program (950-231201), Canada Foundation for Innovation (31390), Canada First Research Excellence Fund ‘Medicine by Design’ (OMNI-2017-01), Ontario Institute for Regenerative Medicine (OMNI-2017-01), and the University of Toronto Faculty of Medicine Deans Fund (to PMG). Finally, we thank Dr. Majid Ebrahimi and Dr. Louise Moyle for reviewing the final manuscript and for providing critical feedback.

## Author contributions

MAB designed and performed the experiments, analyzed and interpreted the data, and prepared the manuscript. ESL, NRI, HvdD, and TF designed experiments, provided OLIG2^+^ cells, provided training on differentiation of OLIG2^+^ cells to motor neurons, interpreted the data, and revised the manuscript. BM and CTN designed and performed electrophysiological experiments, analyzed and interpreted the data, and prepared the manuscript. KT consented patients, obtained research ethics board approval, and transferred tissue biopsies to MAB. BAS, MSS, MZ, and RSA designed experiments, interpreted data, and revised the manuscript. AB provided immortalized human myoblast cell lines, interpreted data, and revised the manuscript. EP consented patients and collected sera from MG patients. HA and HG surgically resected and provided skeletal muscle biopsies, interpreted the data, and revised the manuscript. PMG designed the experiments, supervised the work, interpreted the data, prepared and revised the manuscript.

## Competing financial interests

The authors declare no competing financial interests.

## Materials & Correspondence

Correspondence should be addressed to Penney M Gilbert.

## Supplemental Information

### Link to Supplemental Figures and Movies

https://www.dropbox.com/sh/bf493j67aw5hc73/AACf5fY3ic9tmSaCstPxcnSua?dl=0

